# Alkyl gallates display elicitor activities in tobacco plants

**DOI:** 10.1101/145425

**Authors:** Pascale Goupil, Razik Benouaret, Claire Richard

## Abstract

Alkyl gallates showed elicitor activities on tobacco in both whole plants and cell suspensions. Methyl gallate (MG), ethyl gallate (EG) and *n-*propyl gallate (PG) infiltration into tobacco leaves induced hypersensitive reaction-like lesions and topical production of autofluorescent compounds revealed under UV light. When sprayed on tobacco plants at 5 mM, EG promoted upregulation of defence-related genes such as the antimicrobial *PR1*, β-1,3-glucanase *PR2*, chitinase *PR3* and osmotin *PR5* target genes. Tobacco BY-2 cells challenged with EG underwent cell death in 48 h, significantly reduced in the presence of the protease inhibitor aprotinin. The three alkyl gallates all caused alkalinisation of the BY-2 extracellular medium, whereas gallic acid did not trigger any pH variation. Using EGTA or LaCl_3_, we showed that Ca^2+^ mobilisation occurred in BY-2 cells elicited with EG. Overall, our findings are the first evidence of alkyl gallate elicitor properties with early perception events on plasma membrane, potential hypersensitive reactions and PR-related downstream defence responses in tobacco.

**Highlights:**

– Alkyl gallates elicited defence reactions in tobacco
– Alkyl gallates induced local biochemical changes in tobacco leaves
– Alkyl gallates caused modification of plasma membrane properties
– Ethyl gallate led to defence transcript accumulation and dose-dependent cell death associated with hypersensitive response
– Alkyl gallates are novel elicitor agents well-suited to crop protection schemes.

## Introduction

Open-field crops are always vulnerable to pathogens, and so extensive farming requires constant control of pathogen populations. Synthetic pesticides are commonly used for direct pest control, but need repeated spreading for season-long protection. Pesticide action is mostly selective and requires combinations of active compounds in an integrated pest management approach. A promising alternative that emerged in the 1990s is based primarily on improving the endogenous defence capabilities of plants. When plants are attacked by pathogens, they defend themselves with an arsenal of defence mechanisms. These active plant defence responses comprise three successive events: (i) recognition of the pathogen, (ii) signal transduction, and (iii) execution of the defence programmes with activation of defence genes (Fu and Dong, 2013). The recognition events involve pathogen- or plant-derived signalling entities termed elicitors (Choi and Klessig, 2016). Intracellular signal conversion and transduction include the very early elicitor-induced signalling processes such as protein phosphorylation, influx of Ca^2+^, efflux of H^+^, K^+^ and Cl^-^ leading to extracellular alkalinisation, depolarisation of the plasma membrane, and active oxygen species production, culminating in the hypersensitive response, a localised cell suicide (Yang *et al*., 2011). Ultimately, the pathogen challenge leads to the production of phytoalexins and pathogenesis-related proteins, which prevent pathogen development. Point inoculation of the pathogen, in addition to triggering the local acquired resistance response (LAR), can mobilise systemic acquired resistance (SAR), which alerts and predisposes the tissues to a more intensive response on subsequent infection (Kachroo and Robin, 2013; Fu and Dong, 2013). Salicylic acid (SA) plays a crucial role in the SAR induction processes. SA is involved in the signal transduction systems by activating enzyme expression catalysing biosynthetic reactions to produce defence compounds (Chen *et al*., 2009; Boatwright and Pajerowska-Mukhtar, 2013) and provide protection for plants against pathogens.

As it is pathogenesis that initiates SAR, impact on disease is lower than if the SAR response had been activated before the arrival of the pathogen. The use of plant defence inducers (PDIs) to establish SAR intentionally and preventively in plants are thus currently a promising strategy in phytoprotection. PDIs are organic, inorganic, botanical or synthetic stimulus-triggering factors (elicitors) that mobilise plant defence reactions (Walters *et al*., 2013; Lyon, 2014). By stimulating plant immunity, PDIs provide defence against subsequent infection (Gozzo and Faoro, 2013; Walters *et al*., 2013). To date, only very few substances with high SAR activity have been described (Gozzo and Faoro, 2013), and they mostly combine PDI and pesticide actions, not necessarily a disadvantage since both properties are inimical to pathogens (Dewen *et al*., 2017).

Agricultural applications of plant activators are far from valued (Gozzo and Faoro, 2013). The synthetic agrochemicals now available are SA-like substances that mimic the action of this phytohormone. Of these, 2,6-dichloro isonicotinic acid (INA) and acibenzolar-*S*-methyl (BTH) and their derivatives are the most thoroughly studied, currently marketed as BOOST^®^, Bion^®^ and Actigard™. These resistance activators serve as efficient chemical inducers of SAR, with effectiveness against diverse plant diseases (Bektas and Eulgen, 2015; Dufour *et al*., 2016).

The identification of new chemicals able to induce disease resistance is beneficial not only for elucidating the pathways leading to SAR, but also for developing new plant protection agents. The effectiveness of various elicitors depends on plant taxonomy, and there are many examples where an elicitor is more effective on some plant species than on others (Lyon, 2014). We have been engaged in seeking naturally-occurring or synthetic substances able to induce SAR reactions. As part of this research we evaluated the PDI activity of alkyl gallates (3,4,5-trihydroxybenzoates) on tobacco plants. Gallic acid derivatives had already been found to display antimicrobial properties. Octyl, nonyl and lauryl gallates exhibit fungicidal and bacterial activity (Kubo *et al*., 2002; Leal *et al*., 2009). Silva *et al*. (2013) showed their potential for use in disease management, evaluating the inhibitory activity of a set of gallic acid alkyl esters on the plant pathogenic bacterium *Xanthomonas citri*. Here we demonstrate for the first time that alkyl gallates act at various points in the signalling pathway leading to defence reactions. We report evidence that treatment of tobacco with methyl, ethyl and *n*-propyl gallates induces hypersensitive-like reactions with activation of defence-related genes in the tobacco plant. These gallic acid derivatives trigger rapid extracellular alkalinisation of BY-2 plant cells with Ca^2+^ mobilisation, suggesting that tobacco cells perceive the chemicals by modifying plasma membrane physical properties. Finally, the naturally-occurring and semisynthetic gallic acid derivatives revealed potent PDI actions, revealing a valuable source of compounds that could find use in green agriculture practices.

## Materials and Methods

### Biological compounds and chemicals

Methyl gallate (MG), ethyl gallate (EG), *n-*propyl gallate (PG), salicylic acid (SA), and gallic acid (GA) were purchased from Sigma-Aldrich (Saint-Quentin-Fallavier, France). All the compounds were dissolved in ultrapure water (MilliQ). PG needed warm (25 °C) ultrapure water to dissolve completely.

### Plant material and treatments

The biological activity of alkyl gallates was measured on 3-month old tobacco plants (*Nicotiana tabacum* L.) bearing 20–22 leaves (preflowering stage). Tobacco plants were grown in a greenhouse under controlled conditions (22 ± 5 °C with a photoperiod of 16 h light). The chemical solutions to be tested (50 µl) were infiltrated into the leaf tissue using a plastic syringe (without a needle), on an area of 1–2 cm^2^. As a positive control, leaves were infiltrated with 2 mM SA. For the negative control, leaves were infiltrated with ultrapure water. The compounds were infiltrated into distinct areas on the same leaf for the examination of macroscopic symptoms under bright field or UV light (at 312 nm). For PR transcript quantification, three mature leaves were sprayed on both their upper (adaxial) and lower (abaxial) surfaces. Foliar spray treatments were administered with a fine atomiser until the solutions dripped (3 mL per leaf on average). The negative control was leaves sprayed with ultrapure water.

Tobacco cell suspensions of *Nicotiana tabacum* L. cv. Bright Yellow-2 (BY-2) were grown in MS medium (Duchefa, Netherlands) supplemented with sucrose (30 g.L^-1^), thiamine (1 mg.L^-1^), myo-inositol (102 mg.L^-1^) and 2,4-dichlorophenoxyacetic acid (0.2 mg.L^-1^) at pH 5.8. Cells were agitated in the dark (140 rpm, 25 °C) and subcultured weekly by dilution at a 1:15 ratio in fresh medium. For determination of extracellular pH changes, cells from culture in exponential-phase growth were collected by filtration, washed with 175 mM mannitol, 0.5 mM CaCl_2_, 0.25 mM MgCl_2_, 1 mM KCl and 1 mM sucrose, and resuspended at 0.1 g fresh weight (FW)/mL in the same medium. The initial pH was about 4.8. For transcript accumulation analysis, cells were collected by filtration, and resuspended at 0.3 g FW/mL in fresh MS medium using open 6-well microplates (Greiner Bio-one, Germany). Cells were first equilibrated for 2 h with continuous stirring before elicitation with alkyl gallates.

### Transcription levels of defence genes

The patterns of PR gene expression were analysed by real-time quantitative PCR (RT-qPCR) using SYBR-green and the iCycler iQv5 (Bio-Rad). Total RNA was extracted from tobacco leaf tissue (200 mg) in cetyltrimethylammonium bromide extraction buffer, as described by Chang *et al.* (1993). Total RNA was cleaned up with 0.5 U DNase I solution (Euromedex, France) containing 20 U RNase inhibitor (Euromedex, France). RNA integrity was verified on a 1% agarose gel by detecting ribosomal RNA. RNA concentrations were measured spectrophotometrically at 260 nm. First-strand cDNA was synthesised from up to 1 µg of total RNA with Euroscript Reverse Transcriptase (Eurogentec, France) according to the manufacturer’s instructions. PCR reactions were prepared using the qPCR kit Mastermix for SYBR green (Eurogentec) according to the manufacturer’s protocol. The cDNA concentration used produced a threshold cycle value (*C*_T_) of between 15 and 30 cycles. Amplification specificity was checked by melting-curve analysis. The relative quantity (*Q*_R_) of PR gene transcripts using the EF-1α gene (Peng *et al*., 2004) as internal standard was calculated with the delta-delta mathematical model (Pfaffl 2001). Values are given as log_2_*Q*_R_. For every PR gene analysed, three independent biological replicates were run, and every run was carried out at least in triplicate. Primers and amplicon sizes are given in Benouaret and Goupil (2015).

### BY-2 Extracellular alkalinisation

The pH variations of the culture medium were recorded by introducing a glass microelectrode (Hanna Instrument, HI1330B, France) into 6 ml of the equilibrated cell suspension culture. The pH was measured every 10 min starting immediately after treatment with elicitors or ultrapure water (negative control). The ΔpH was the pH measured every 10 min relative to the pH measured at *t*_0_ immediately after adding elicitors or ultrapure water. Each experiment was repeated at least three times to check reproducibility. Figures depict the results of typical experiments. Pharmacological treatments were conducted using the chelator EGTA (2 mM) and the Ca^2+^ channel blocker LaCl_3_ (2 mM).

### Cell death measurement

The Evans Blue assay was performed according to Amano *et al.* (2003) using a plastic syringe device. The pigment enters cells with damaged plasmalemma. The assay was first calibrated with a set of viability standards using mixtures of living and dead tobacco BY-2 cells (Benouaret *et al*., 2014). Tobacco cells (500 µL) were withdrawn from the culture medium immediately (*t*_0_) and then 24 h after elicitation. Pharmacological treatment was conducted with the serine protease inhibitor aprotinin (0.2 µM) 30 min before cell death rate determination. Absorbance at 600 nm was measured to estimate cell death. Each experiment was repeated three times with three independent BY-2 cultures.

## Results

### Macroscopic changes of tobacco tissues challenged with alkyl gallates

Foliar tobacco tissues were infiltrated with the three alkyl gallates (Fig. 1) at 5 mM, and the macroscopic changes were examined under bright and UV light. Fig. 2 shows the kinetics of appearance of macroscopic symptoms on the abaxial face of tobacco leaves exposed to bright light (Fig. 2A-D) and UV light (Fig. 2E–H). UV light examination (*λ* = 312 nm) revealed that the methyl gallate (MG)-, ethyl gallate (EG)- and *n*-propyl gallate (PG)-infiltrated zones and the surrounding tissues displayed autofluorescence indicative of the accumulation of phenolic compounds (Fig. 2F–H). At 2 and 4 dpi, tobacco mesophyll infiltrated with MG was unchanged under bright light, but the chemical induced autofluorescence in the surrounding tissues. A restricted chlorotic zone appeared at 8 dpi without production of fluorescent compounds. By contrast, the EG-infiltrated tissues turned chlorotic by 4 dpi and tissue injury was enlarged at 8 dpi. The surrounding zone was loaded with phenolic compounds. The PG-infiltrated tissues showed the most pronounced signs, with restricted brown areas at 2 dpi and a brown extended zone at 4 and 8 dpi. Examination of the leaf tissue by UV light revealed that the necrotic zone was surrounded by a scattered pattern of fluorescence. Overall, tobacco leaves were receptive to the three alkyl gallates tested. Since the alkyl gallates themselves did not emit fluorescence (Fig. 2E), these results evidence that the chemical compounds induced marked metabolic changes over the interveinal areas of the leaves. The naturally-occurring EG was retained for further work, since it proved highly active in inducing accumulation of autofluorescent compounds.

**Fig. 1.**
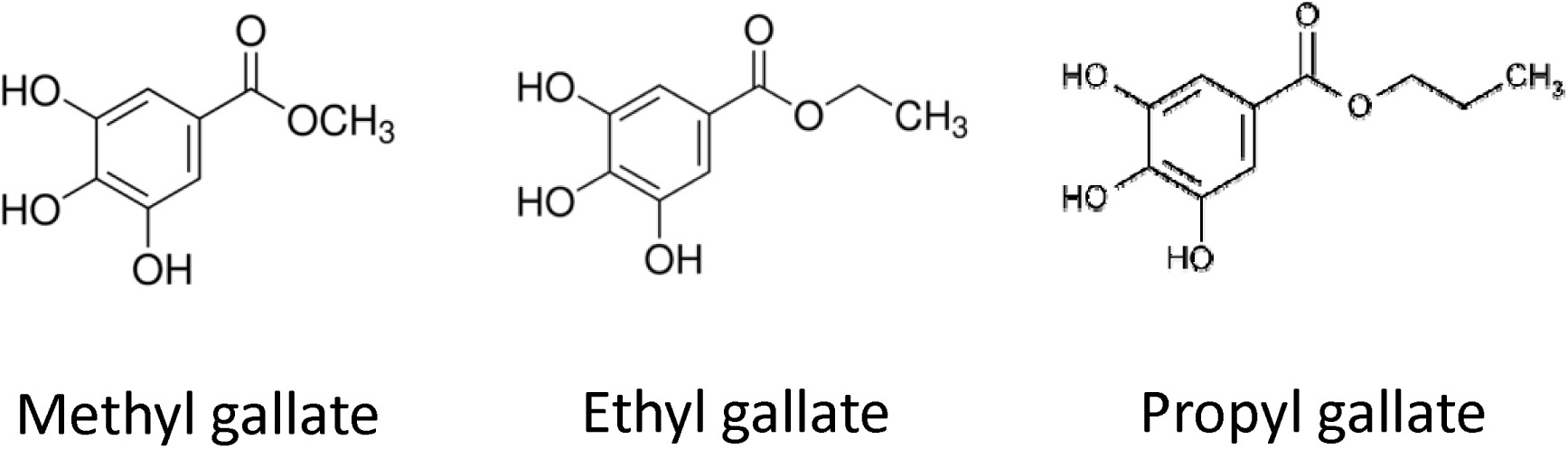
Structures of the active ingredients studied

**Fig. 2.**
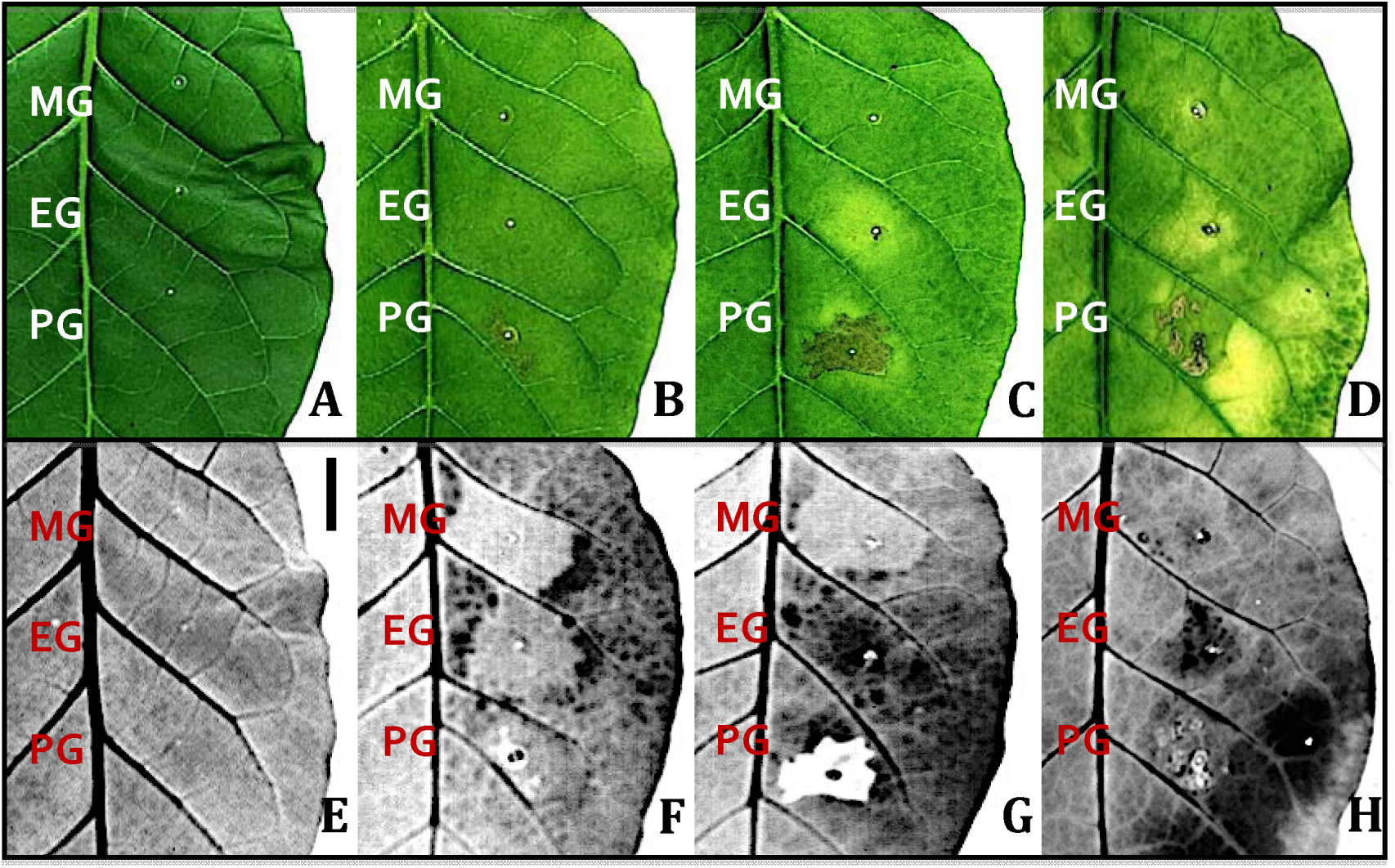
Macroscopic symptoms induced in tobacco leaves by methyl gallate (MG), ethyl gallate (EG) and *n-*propyl gallate (PG) infiltration observed under bright field (A-D) and UV light (E-H) at 0 dpi (A,E), 2 dpi (B,F), 4 dpi (C,G) and 8 dpi (D,H). Alkyl gallates were infiltrated at 5 mM. Bars 2 cm

A dose-response experiment was undertaken on EG-infiltrated tobacco tissues. Macroscopic symptoms were examined under bright and UV light for EG concentrations ranging from 5 mM to 0.5 mM. Fig. 3 shows the infiltrated areas at 4 dpi. The 5 mM-infiltrated zone appeared slightly chlorotic and displayed a large fluorescent surrounding area. Expansion of autofluorescence was induced dose-dependently. EG infiltration at 2 mM resulted in faint autofluorescence, and at 1 mM had no effect. As expected, SA (2 mM) induced a necrotic area with local accumulation of fluorescent compounds.

**Fig. 3.**
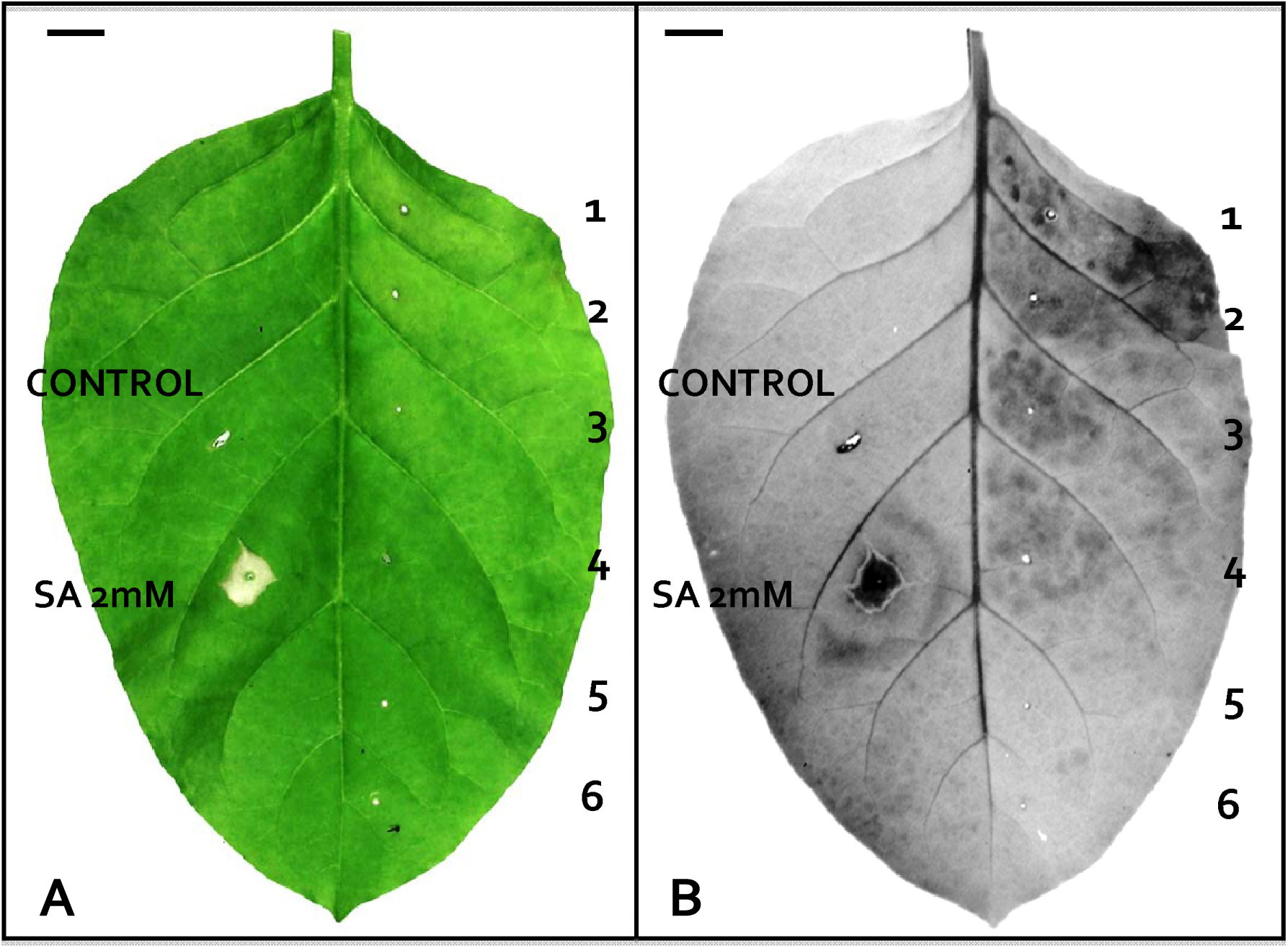
Macroscopic symptoms induced in tobacco leaves by EG infiltration observed under bright field (A) and UV light (B) at 4 dpi. Tobacco leaves were infiltrated with different EG concentrations: 5 mM (1), 4 mM (2), 3 mM (3), 2 mM (4), 1 mM (5) and 0.5 mM (6), ultrapure water (control), 2 mM SA. Bars 1 cm

### Responses of cultured cells to alkyl gallates

We used medium alkalinisation as a readily measurable effect on suspension-cultured BY-2 cells to test the potential elicitor activity of alkyl gallates. As shown in Fig. 4A, alkalinisation of the culture medium in response to AG was evidenced for the 1 mM and 5 mM concentrations. Suspended cells responded to AG by a rapid, sustained alkalinisation of the incubation medium. Extracellular alkalinisation was detected immediately after adding the AG. The three alkyl gallates induced a ΔpH reaching 0.8 to 1.1 units at 1 mM and 1.5 to 1.8 units at 5 mM after 150 min. Fig. 4B shows that the extent of pH variation was dose-dependent in the concentration range used for EG. The threshold value was not reached in our experimental system; the lowest concentration (0.5 mM) could still cause a pH change of 0.8 units. As expected, no pH shift was detected with the control cells treated with water.

**Fig. 4.**
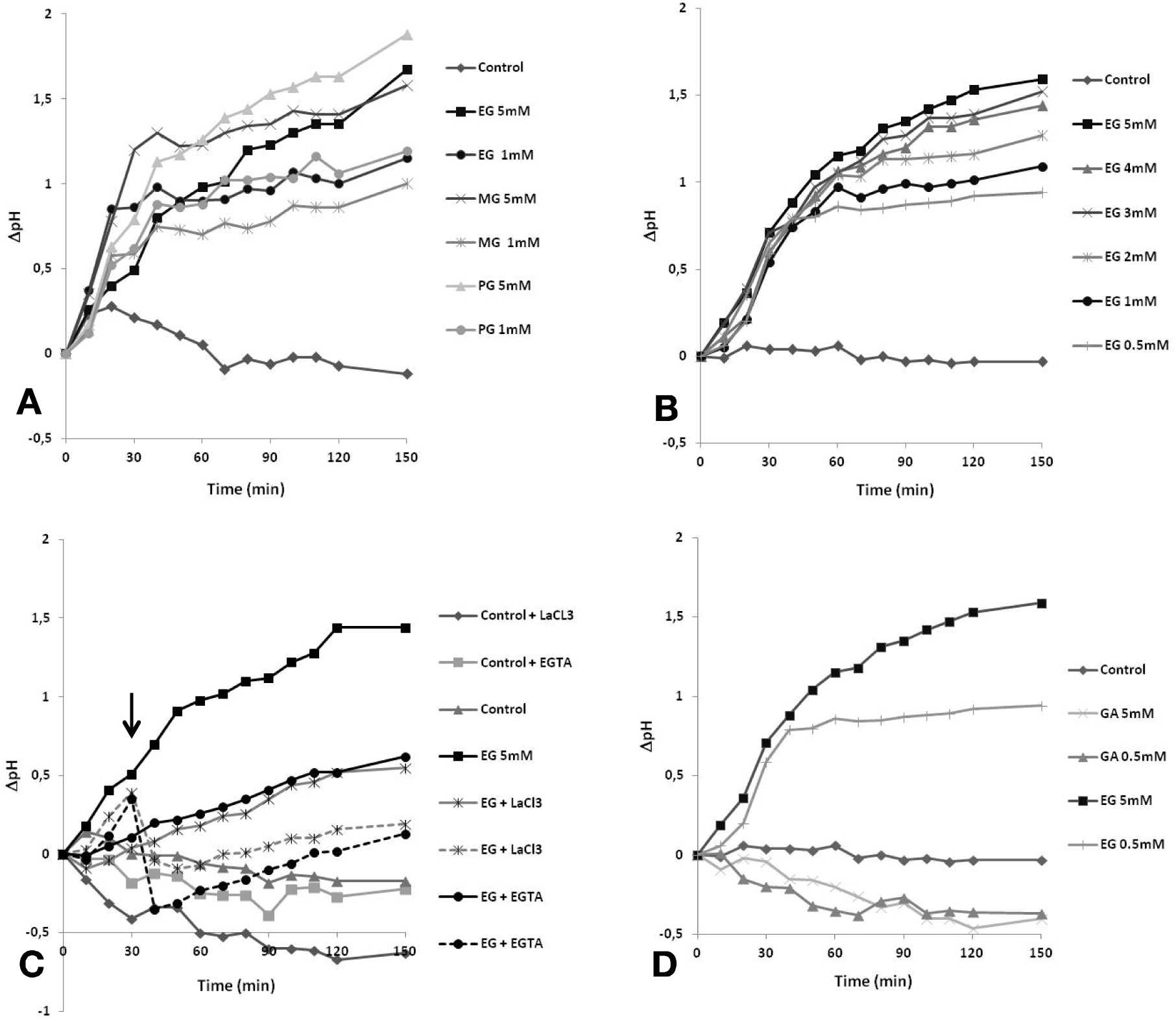
Extracellular alkalinisation in tobacco BY-2 suspension cells over time elicited with methyl gallate (MG), ethyl gallate (EG), and *n-*propyl gallate (PG). A. Alkyl gallates used at 5 and 1 mM, B. EG in the presence of Ca^2+^ inhibitors. EGTA and LaCl_3_ at 2 mM were added to the BY-2 medium at the onset of elicitation or at 30 min (arrow), C. EG added at 0.5 to 5 mM, D. Gallic acid (GA) and EG used at 5 and 0.5 mM. The control is water. The data are from representative experiments out of a total of five.

The calcium mobilisation in EG-induced pH changes was investigated by adding the chelator agent EGTA (2 mM) and the Ca^2+^-channel blocker LaCl_3_ (2 mM) to the culture medium concomitantly with the EG. Both chemicals radically suppressed the pH shift. Adding the chemicals during the alkalinisation phase, i.e. 30 min after elicitation, resulted in rapid abolition of the pH rise. EDTA and LaCl_3_ had an inverse effect on the extracellular pH medium in control cells, with slight medium acidification (Fig. 4C).

The eliciting activity of gallic acid (GA) was measured by medium pH determination. Addition of GA at 0.5 mM and 5 mM to the BY-2 cell suspension did not trigger any pH variation in the extracellular medium, suggesting that the compound was not perceived by the tobacco cells, at least as noted above (Fig. 4D).

The EG-triggered hypersensitive-like response was investigated in BY-2 cells using the Evans blue test (Fig. 5). Cell death rate was measured in tobacco cells exposed to EG at 0.5 mM, 2 mM and 5 mM for 48 h (Fig. 5A). Cell death rate was 10% on average in control cells (without EG). The EG elicitation induced significantly higher cell death. The exposure of tobacco cells to 2 mM and 5 mM EG increased the magnitude of the cell death rate, which reached 25% and 45% respectively. The action of the plant cell death inhibitor aprotinin was examined on 5 mM EG-elicited cells. As shown in Fig. 5B, the protease inhibitor lowered cell death rate by 19%, whereas the inhibitor did not affect cell viability in control tobacco cells.

**Fig. 5.**
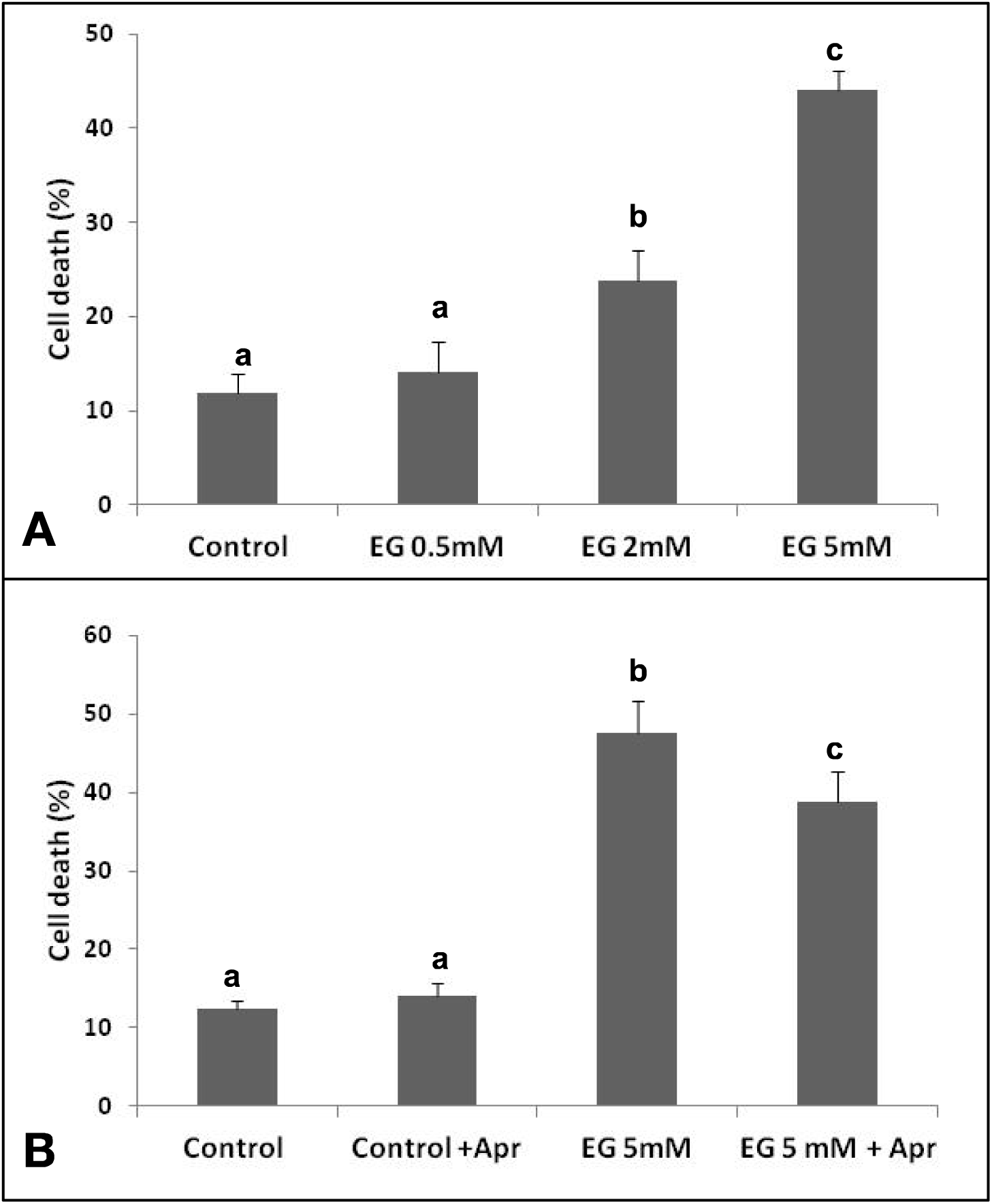
Cell death rate assessed by Evans Blue staining of tobacco BY-2 cell suspension treated with ethyl gallate (EG) at 0.5, 2 and 5 mM for 48 h. The protease inhibitor aprotinin was added 30 min before cell death measurement in BY-2 cells treated with water (control) or 5 mM EG. Values are expressed relative to control (water treatment) values. The experiments were performed in triplicate with three different BY-2 cultures. Bars represent means ±SEM. Letters indicate significant differences between treated and control samples according to one-way ANOVA followed by LSD test (*p* ≤ 0.05). Bars represent the mean values ± SE, *p* ≤ 0.05.

### Ethyl gallate-induced PR transcript accumulation

The ability of EG to induce defence-related gene expression was investigated in tobacco plants. Transcript levels of four PR target genes were quantified by quantitative real-time PCR in EG-treated leaves 4 days after spraying. At 1 mM, EG failed to induce significant transcript accumulation of the four PR markers. At 5 mM, EG led to high PR transcript accumulation: 60-fold for PR1, 50-fold for PR2, 35-fold for PR3, and 30-fold for PR5, on average (Fig. 6).

**Fig. 6.**
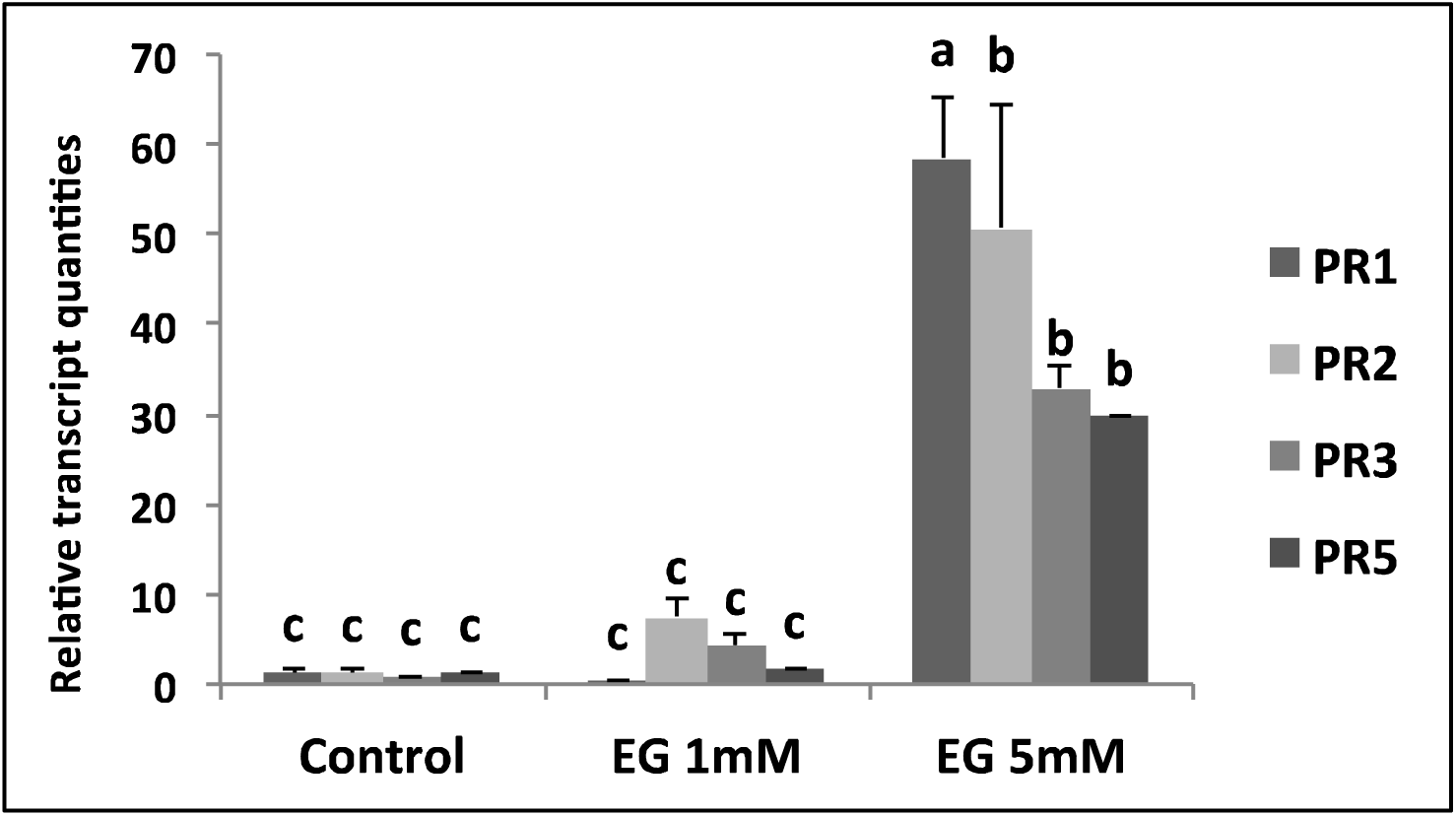
PR transcript accumulation in tobacco leaves 4 days after 2 mM SA (positive control) or 1 and 5 mM ethyl gallate (EG) spraying. Transcripts were quantified by real-time RT-PCR in the sprayed leaves. Values are expressed relative to control (water treatment) values. Three leaves per plant were treated for each experiment. Each independent experiment was performed in triplicate. Bars represent means ± SEM. Letters indicate significant differences between treated and control samples according to one-way ANOVA followed by LSD test (*p* ≤ 0.05).

## Discussion

The present work demonstrates the ability of three alkyl gallates to elicit plant defence reactions. Using tobacco as a plant model, we show the potential of methyl gallate (MG), ethyl gallate (EG) and *n*-propyl gallate (PG) to trigger local injuries, biochemical changes and PR transcript accumulation. Infiltration of tobacco leaves with the chemicals resulted in foliar symptoms and local accumulation of fluorescent compounds. The three compounds induced restricted chlorotic tissues visualised under bright light, with production of phenolics in the surrounding areas, suggesting cell death initiation of the mesophyll cells. The topical reactions induced by the three alkyl gallates were of different severity in terms of tissue injury and production of phytoalexin-like entities. The fact that the intensity of effects varied with the alkyl gallates (MG < EG < PG) suggests that the compounds were active to different degrees at the 5 mM concentration, or that their modes of action differed in the necrotic reaction associated with cell death. EG and PG seemed to be the most potent inducers of local cell death in our experiment. Localised cell death in plant tissue resulting from attack by pathogens can occur either in hypersensitive resistant response or in harmful reactions such as necrosis potentially caused by toxins (Iakimova *et al*., 2005). We demonstrate here that BY-2 cell death is sensitive to protease inhibitor, suggesting that infiltrated tobacco tissues may evolve towards hypersensitive status. However, phytotoxic reactions caused by alkyl gallate cannot be ruled out.

Rapid changes in membrane permeability are involved in plant defence-associated early responses to various biotic elicitors (Garcia-Brugger *et al*., 2006). Increased influx of protons and calcium mobilisation are both early events in the elicitor signalling network. Perception of alkyl gallates visualised by the pH rise in the incubation medium was evidenced on BY-2 cells. Medium alkalinisation is one of the very early events occurring in elicitor-binding sites in plant cells (Zhao *et al*., 2005), and results from elicitor-induced depolarisation of the plasma membrane with subsequent K^+^/H^+^ exchanges and Ca^2+^ influx/Cl^-^ efflux. In our experiments, the effect of Ca^2+^ influx inhibitors EGTA or LaCl_3_ caused the decreased pH shift of the extracellular medium, suggesting that Ca^2+^ mobilisation occurred in BY-2 cells elicited with alkyl gallates. As a consequence of ion fluxes, a predictive cytoplasmic acidification should take place rapidly in BY-2 cells. Cytoplasmic acidification is regarded as an essential step in signal transduction, leading to early defence gene expression and secondary metabolite biosynthesis (Sakano, 2001).

Kinetics of extracellular pH changes vary among elicitors, with two main different behaviours towards cells. According to the nature of the elicitor, the perception can induce a long-sustained pH rise, or the pH variation can be transient, with a return to the resting value. It is noteworthy that the modification of H^+^ flux in BY-2 cells following treatment by alkyl gallates was similar to the defence response elicited by a polyphenolic-enriched grape extract (Benouaret *et al*., 2014). The induced H^+^ flux modification lasting 150 min presented similar characteristics in the respective time course of the effect. This result suggests that the underlying processes are basically the same. EG is a naturally-occurring gallic acid derivative found in several plant sources and identified in wines from different varieties of *Vitis vinifera* (Monagas *et al*., 2005). It was not detected in a red winery byproduct extract (data not shown), suggesting that this biomolecule is not active in grape marc extract.

Alkyl gallates are antioxidants. One of the earliest cellular responses following elicitor recognition is oxidative burst with production of active oxygen species (AOS). Despite their damaging effect, AOS are well-described second messengers producing plant responses. Whether AOS will act as a destructor or as a signalling entity depends on the equilibrium between AOS production and scavenging. Low concentrations of AOS act as signalling entities mediating the plant responses: however, high concentrations cause acute damage to cell components (Sharma *et al*., 2012). In elicitor-challenged cells, disruption in cellular redox homeostasis appear to switch on a signalling cascade leading to programmed cell death, encountered during hypersensitive response (HR) (Van Breusegem and Dat, 2006). Using pharmacological approaches, our group came near to the hypersensitive status of BY-2 cells that led to cell death on EG treatment. In this BY-2 system, the antioxidant gallates acting as free radical scavengers could subsequently help to reduce physiological AOS equilibrium and induce HR-related cell death. The potential ability of phenolics to induce HR-related cell death has been previously demonstrated with polyphenolic-enriched grape marc extracts (Benouaret *et al*., 2014). Even though EG was not detected in the grape marc extracts, it can be assumed that the antioxidant properties of polyphenolics could be involved in HR establishment. Investigation of AOS as potential agents with pharmacological approaches will help to characterise the pathways leading to the hypersensitive status of EG-induced cell death in tobacco.

Alkyl gallates evidently activate the SAR pathway, since they induced expression of the same set of SAR genes, i.e. *PR1*, *PR2*, *PR3* and *PR5*, that are induced by SA (Cordelier *et al*., 2003). Numerous synthetic and/or bio-based elicitors can induce plant immune responses. The most marked chemical ones, acibenzolar-*S*-methyl (ASM) and 2,6-dichloroisonicotinic acid (INA), both induce SAR without SA accumulation; probenazole (PBZ) induces SAR through SA biosynthesis (Bektas *et al*., 2015; Burketova *et al*., 2015). SA involvement in alkyl gallate-regulated plant defences and the potential of these compounds to protect cultivated crops are being investigated in laboratory studies.

In conclusion, the present work offers evidence for plant immunity inducer properties of alkyl gallates. Plants can synthesise a wide range of phenolics as an integral part of their non-host resistance strategy. These secondary metabolites are natural defensive antibiotics, ensuring efficient chemical barriers to pathogens. Their elicitation properties are demonstrated here for the first time in alkyl gallates, which emerge as PDI agents suited to phytoprotection. Being naturally-occurring and ubiquitous, EG is the most promising of these agents. It is potentially non-toxic for the human being, and offers consumer benefits, with antioxidant, anti-inflammatory and anticarcinogenic activities. Given that EG can play a dual role in plant protection, as it also possesses direct antimicrobial activities, this biocontrol agent constitutes a potent product well-suited to environmentally-oriented schemes for high crop protection.

## Acknowledgements

The authors thank Céline Sac for help with the tobacco cell cultures and laboratory assistance, and Dominique Marcon for technical assistance in photographic editing.

